# Using Piecewise Regression to Identify Biological Phenomena in Biotelemetry Datasets

**DOI:** 10.1101/2021.12.14.472652

**Authors:** David W. Wolfson, David E. Andersen, John R. Fieberg

**Affiliations:** University of Minnesota, Minnesota Cooperative Fish and Wildlife Research Unit; U.S. Geological Survey, Minnesota Cooperative Fish and Wildlife Research Unit; University of Minnesota

## Abstract

Technological advances in the field of animal tracking have greatly expanded the potential to remotely monitor animals, opening the door to exploring how animals shift their behavior over time or respond to external stimuli. A wide variety of animal-borne sensors can provide information on an animal’s location, movement characteristics, external environmental conditions, and internal physiological status. Here, we demonstrate how piecewise regression can be used to identify the presence and timing of potential shifts in a variety of biological responses using GPS telemetry and other biologging data streams. Different biological latent states can be inferred by partitioning a time-series into multiple segments based on changes in modeled responses (e.g., their mean, variance, trend, degree of autocorrelation) and specifying a unique model structure for each interval. We provide five example applications highlighting a variety of taxonomic species, data streams, timescales, and biological phenomena. These examples include a short-term behavioral response (flee and return) by a trumpeter swan (*Cygnus buccinator)* immediately following a GPS collar deployment; remote identification of parturition based on movements by a pregnant moose (*Alces alces*); a physiological response (spike in heart-rate) in a black bear (*Ursus americanus*) to a stressful stimulus (presence of a drone); a mortality event of a trumpeter swan signaled by changes in collar temperature and Overall Dynamic Body Acceleration; and an unsupervised method for identifying the onset, return, duration, and staging use of sandhill crane (*Antigone canadensis*) migration. We implement analyses using the mcp package in R, which provides functionality for specifying and fitting a wide variety of user-defined model structures in a Bayesian framework and methods for assessing and comparing models using information criterion and cross-validation measures. This approach uses simple modeling approaches that are accessible to a wide audience and is a straightforward means of assessing a variety of biologically relevant changes in animal behavior.

## Introduction

Recent technological advancements in the field of biotelemetry have greatly expanded the potential for remotely monitoring animals (Cagnacci et al., 2010; Hebblewhite & Haydon, 2010; Tomkiewicz et al., 2010). Historically, animals were tracked using very-high frequency (VHF) telemetry, which requires a receiver in close proximity to an animal to triangulate its location (Craighead, 1982). Global positioning system (GPS) transmitters, now the standard for most wildlife studies, are small enough to be placed on bats and songbirds weighing less than 20 grams (Cvikel et al., 2015). Current GPS technology provides copious fine-scale data over long study durations and spatial coverages. Other biologging sensors such as accelerometers, temperature-depth recorders, geolocators, and even heart-rate monitors, also produce valuable, high-frequency data on physiological and external conditions affecting animals (Cooke et al., 2004; Jonsen et al., 2007; Wilmers et al., 2015). Such data can elucidate behavioral patterns (foraging, migration, predator/prey dynamics) and provide knowledge about demographic parameters such as fecundity and survival; however, processing high-volume data can be complex and time-consuming (Brooks et al., 2019; Hamilton et al., 2017; Kramer et al., 2018). Simple and flexible tools are necessary to facilitate efficient analysis of animal biotelemetry data to examine behavior over time and response to external stimuli.

When inferring behavior, raw measurements (e.g. location, acceleration, body temperature) can be used, or these data may be first converted into metrics that provide additional insights into biological events or phenomena (e.g. net-squared displacement for migration, overall dynamic body acceleration for energy expenditure: Wilson et al., 2006; Bunnefeld et al., 2011). Data collected at a high frequency provide autocorrelated time series, which can preclude some modeling approaches, such as resource selection functions, that require observations to be statistically independent. Past approaches for accounting for autocorrelation have focused on filtering data, which results in potential loss of information. More recent approaches now directly model the correlation structure to provide additional inference without loss of information (Fleming et al., 2015, Fleming et al, 2016).

Analyses of biotelemetry data collected at a high frequency often begin by partitioning datasets into homogeneous segments that correspond to different behavioral states (Edelhoff et al., 2016). Some common methods for partitioning datasets include clustering algorithms that minimize a cost function associated with statistical properties of a time series (e.g. the pruned exact linear time (PELT) algorithm or the penalized contrasts method; Lavielle 2005; Killick et al., 2012) or parametric state-space models such as hidden Markov models (Patterson et al., 2008). Often, these methods require restrictive assumptions, are computationally intensive, or are restricted to a single type of data stream (Morelle et al., 2017). Here, we demonstrate how piecewise regression can be used to analyze a wide range of biotelemetry data sets, offering a flexible and user-friendly approach with results that are easy to interpret.

### Overview of Piecewise Regression and applications with the mcp package

Piecewise regression is a common statistical method used to model ecological thresholds (Toms & Lesperance, 2003) and can be used to identify change points that signify potential shifts in the relationship between response and explanatory variables (Muggeo, 2003; Toms & Villard, 2015). Piecewise regression is an extremely flexible modeling framework due to the ability to specify unique model structures for each segment between change points. Transitions between segments can be abrupt, if segments are disjunct, or smooth, if segments are joined.

Herein, we focus on a Bayesian formulation of piecewise regression following the formulation of Stephens (1994). Let **y** = (*y*_1_, …, *y*_*n*_) be the realization of a sequence of random variables *Y* = (*Y*_1_, …, *Y*_*n*_) of length *n*, with a change point at *r*(1 ≤ *r* ≤ *n*). We can write the distribution of *Y*_1_, …, *Y*_*n*_ as

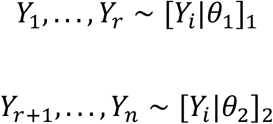

where *θ*_1_ and *θ*_2_ are parameter vectors describing the data-generating process before and after the change point, respectively. Values of *θ* can describe various characteristics of the response distributions, including their means, variances, or correlation structures.

Inference is made via the posterior distribution of *r, θ*_1_, and *θ*_2_, denoted by [*r, θ*_1_, *θ*_2_|*Y*]. From Bayes’s theorem,

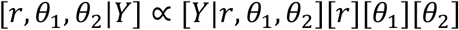

where [*θ*_1_],[*θ*_2_], and [*r*] are prior distributions for the parameters describing the data-generating process. This approach easily extends to multiple change points, *r*_1_, *r*_2_, …, *r*_*k*_ with unique parameter vectors, *θ*_1_, …, *θ*_*k*_, describing the likelihood of *Y* within each segment:

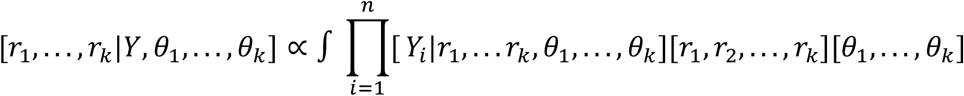

First, consider a simple example represented by separate intercepts before and after a single change point. Let *r* be the location of the change point along the x-axis, where the expected value of *y*_*i*_, *µ*_*i*_, changes from *f*_1_(*x, β*_1_), to *f*_2_(*x, β*_2_):

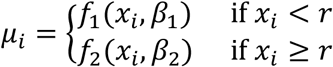

*f*_1_(*x, β*_1_) and *f*_2_(*x, β*_2_) can represent combinations of either additive or multiplicative effects of covariates on segments using the generalized format *f*_*k*_(*x*_*i*_, *β*_*k*_) = *β*_*k*,0_ + *x*_*i*_ · *β*_*k*,1_+…*x*_*i*_ · *β*_*k,p*_ or *f*_*k*_(*x*_*i*_, *β*_*k*_) = *β*_*k*,0_ · *x*_*i*_ · *β*_*k*,1_ ·…*x*_*i*_ · *β*_*k,p*_ with parameters (*β*_*k*,0_, *β*_*k*,1_, …, *β*_*k,p*_) and *k* segments resulting from *k* − 1 change points. For a more detailed description see Lindeløv (2020).

The mcp package uses indicator functions to map elements of a dataset (i.e., segments) to statistical models specific to each segment. For a dataset with y as the response variable and x as the predictor variable, the two-intercept case can be written as:

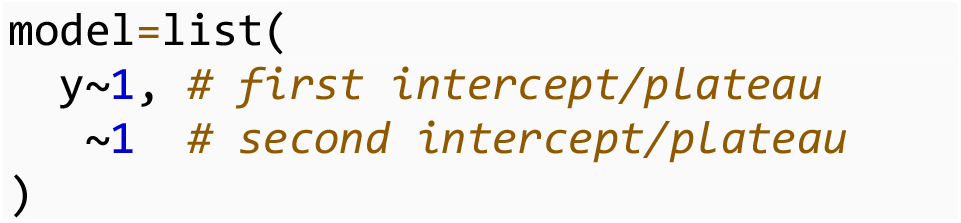

Specifying a model structure with a joined segment is also straightforward:

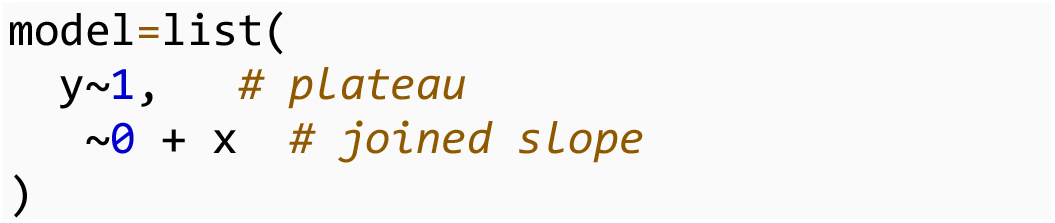

The ‘∼0’ term specifies that the second segment shares the same intercept, whereas the ‘+ x’ term assigns a slope term associated with the variable *x* such that the slope is 0 before the change point and *β* · *x* afterwards.

The mcp package makes it easy to specify custom model structures for each segment and allows for a wide range of statistical distributions and link functions. Model parameters are assigned suitable vague prior distributions, though these can easily be altered (Lindeløv, 2020). Hierarchical models with random effects can be specified by allowing change points to covary within a group, thereby allowing the estimation of individual-specific and population-level change point parameters.

Another advantage of the mcp package is the combination of a user-friendly interface connected to a robust Bayesian modeling backend. The model structure for each segment is expressed using the familiar syntax of the lme4 and brms packages, then is converted ‘on-the-fly’ to JAGS code (Bates et al., 2014; Bürkner, 2017). Support for fitting models using Stan is currently in development. The mcp package returns samples from the marginal posterior distributions for all change points and other model parameters, which can be used to visualize model output, for example with the ggplot2 and bayesplot packages and produce credible intervals (Gabry, 2017; Gabry et al., 2019; Wickham, 2011).

Lastly, the mcp package allows for a robust suite of methods for model assessment and comparison, hypothesis testing, and data simulation. Markov-chain Monte Carlo (MCMC) sampler performance can be checked via the Gelman-Rubin statistic, effective sample size, and visual assessment of MCMC chains. Posterior predictive checks ensure that the fitted model is consistent with the data-generating process (i.e., simulations from the fitted model resemble the original data; Gelman et al., 1996; Gelman & Shalizi, 2013). The influence of the priors can be evaluated by comparing prior-posterior overlap (Youngflesh, 2018). The predictive performance of multiple models can be compared with leave-one-out cross validation (LOO-CV), either using WAIC (Widely Applicable Information Criterion) or ELPD (Expected Log Predictive Density), which can be calculated with the loo package (Gelman et al., 2014; Vehtari et al., 2017). If the appropriate number of change points is not known *a priori*, LOO-CV allows the user to compare the predictive performance of multiple models to determine the optimal number, or to compare different model structures and prior distributions. Additionally, hypothesis testing can be performed using point Bayes factors (i.e., the prior to posterior ratios associated with specific parameter values; Verdinelli & Wasserman, 1995; Wagenmakers et al., 2010). Bayes factors can also be used to compare models that represent alternative hypotheses and for model averaging (Hooten & Hobbs, 2015; Kass & Raftery, 1995). Bayes factors > 10 are commonly used to indicate strong evidence that the data support one model over another (De Santis, 2004; Schönbrodt & Wagenmakers, 2018).

## Example Applications

We demonstrate the use of piecewise regression using the mcp package with five example applications highlighting a variety of taxonomic species, data streams, timescales, and biological phenomena. We have provided R code associated with each example at the publicly available Data Repository for University of Minnesota.

### Identification of altered behavior post-capture

Effects of capture, handling, and transmitter deployment are not well understood for many wildlife species, especially in the short-term period immediately post-capture. Efforts to quantify animal response have predominantly focused on the impacts of the transmitter itself (i.e., the weight and aerodynamics of the unit, usually in birds: Evans et al., 2020) or on the physiological effects from chemical immobilization (typically in large mammals: Barron et al., 2010; Brivio et al., 2015; Thompson et al., 2020). Most researchers have sought to quantify effects on vital rates, such as survival and fecundity (Casas et al., 2015; DelGiudice et al., 2005; Lameris & Kleyheeg, 2017), or short-term ethological responses such as changes in the time spent grooming (Kölzsch et al., 2016; Rachlow et al., 2014). It is generally accepted that events such as capture and handling may result in a short-term period of altered movement behavior. Thus, it is common to remove data from the first week or two post-capture, though often without biological or empirical justification for the threshold used to filter the data.

Arbitrarily filtering data without knowing the existence and duration of capture effects presents several issues for movement-related analyses. Removing data potentially discards useful information, which may already be scarce in studies using VHF telemetry or with small sample sizes (Girard et al., 2002). Alternatively, quantifying behavioral responses post-capture can provide species-specific information on altered behavior and inform future studies (Dechen Quinn et al., 2012). For example, Stabach et al. (2020) examined effects of GPS collar deployment on scimitar-horned oryx (*Oryx dammah*) to quantify the short-term responses in activity, behavior, and stress levels, and the length of time before these effects subsided. Visual observation showed headshaking significantly increased post-capture but returned to pre-capture levels within 3 days, and random forest classification of tri-axial accelerometer data indicated a 480% increase in headshaking compared to stable baseline levels that resumed after 24 hours. Using piecewise regression, Stabach et al. (2020) found that fecal glucocorticoid metabolite levels were elevated for five days following collar deployment, suggesting a stress response, before returning to baseline levels.

The effects of capture and handling on movement may also provide insights into how individuals respond to risky situations (e.g., capture, predation). For example, animals may exhibit a “flee and return” movement response, the strength of which may be indicative of their tolerance of risk as it relates to a fecundity-survival trade-off (Ghalambor & Martin, 2001; Montgomerie & Weatherhead, 1988). DelGiudice et al. (2015) documented that capture of moose (*Alces alces*) neonates caused some mothers to abandon their calves, and Obermoller et al. (2019) found that adult female moose frequently fled when their calves were predated and then returned after the risk had subsided. These responses suggest that many adult female moose favor individual survival over protection of their young.

Other studies have found similar “flee and return” movements in response to human activity such as hunting or helicopter-based capture (Jung et al., 2019; Sunde et al., 2009; Thurfjell et al., 2013). In an ongoing study of trumpeter swan (*Cygnus buccinator*) movement ecology, we found some swans exhibited a distinctive “flee and return” response immediately post-capture (Wolfson unpublished data). Figure 1 shows the results of a piecewise regression model fit to the relationship between Net-Squared Displacement (NSD) and time since capture:

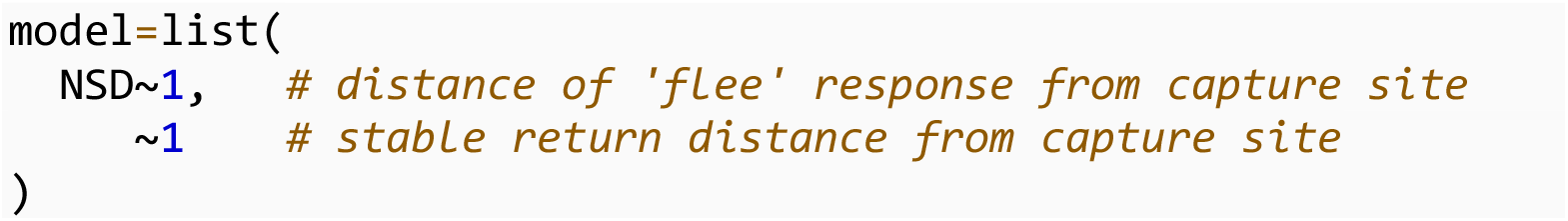

**Figure 1:**
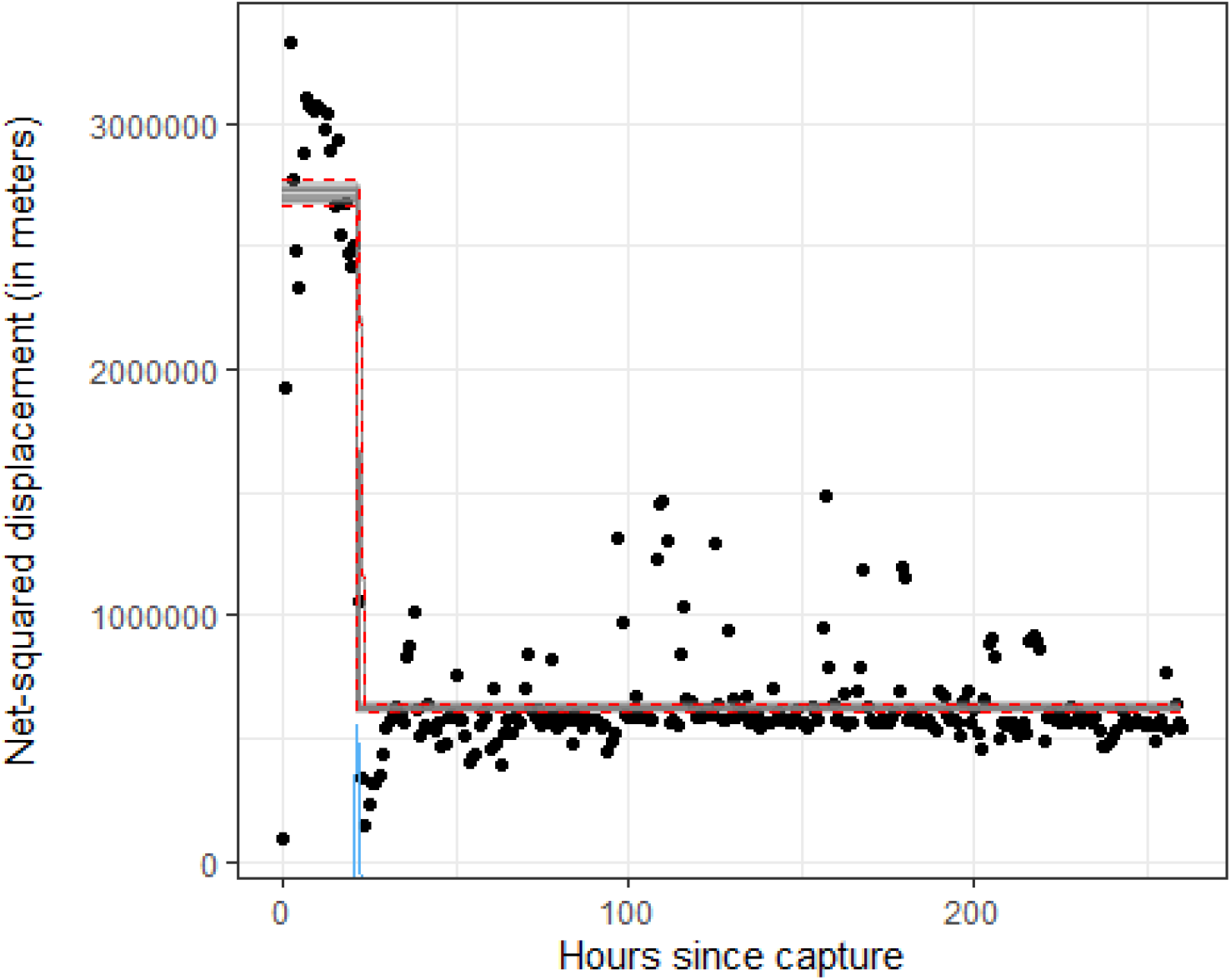
Hourly net-squared displacement measured from the point of release of a Trumpeter Swan (*Cygnus buccinator*) after collar deployment. Grey lines show 25 draws from the posterior distribution, with 95% credible intervals for the mean response shown as red dotted lines. The posterior distribution for the change point is shown in blue on the x-axis.

The NSD value of the first intercept quantifies how far the swan fled and the change point reveals when a return to the capture site occurred. This model, which has two segments with different intercepts and a single change point, had a better fit using LOO-CV to estimate ELPD, a goodness-of-fit measure that estimates the predictive accuracy of a model while balancing the trade-offs of bias and variance, than an alternate model with a single intercept, therefore suggesting the presence of an altered state representing a flee response (Table 1).

**Table 1:** Leave-one-out Cross Validation was used to compute the Estimated Log Predictive Density (ELPD) of two different models; one fit with a single intercept and the other with two intercepts separated by a changepoint. The higher ELPD for the model with two intercepts indicates that it has a higher predictive accuracy.

| Model Syntax | ELPD Difference | SE of Difference |
| --- | --- | --- |
| Two Intercepts | 0.0 | 0.0 |
| One Intercept | -213.3 | 64.2 |

### Pre-parturition movement

Accurate estimation of vital rates and their relative contributions to population dynamics is an essential tenet for population management (Coulson et al., 2005). Signals in movement data can reveal important biological events such as parturition in large ungulates, which relate to vital rates. Although fecundity is a key vital rate, detection of the occurrence and timing of parturition can be difficult, especially in ‘hider’ species that have low parental care of neonates (Lent, 1974; Ralls et al., 1986).

Pregnant ungulates typically make a long-distance movement immediately preceding parturition and then remain sedentary for some period (McGraw et al., 2014). Patterns in movement data thus offer a cost-effective means for identifying parturition in radio-collared ungulates (Asher et al., 2014; DeMars et al., 2013; Nicholson et al., 2019; Peterson et al., 2018; Severud et al., 2015). Although these methods can be effective at inferring parturition events, many are sensitive to parameter choices or involve visual observation of movement metrics, which can be time-intensive (Dettki & Ericsson, 2008).

To demonstrate remote identification of a confirmed parturition event, we provide an example of displacement of a pregnant moose (Figure 2). Severud et al. (2015) confirmed the parturition period by locating and collaring the twin calves shortly after they were born. In this case, the first segment is modeled with an intercept and slope of zero, representing typical moose movement, and the second segment is modeled with a change in the mean displacement (i.e., a different intercept) and also a change in variance (“sigma” in code below) that is representative of a shift to more sedentary movement at time of parturition.

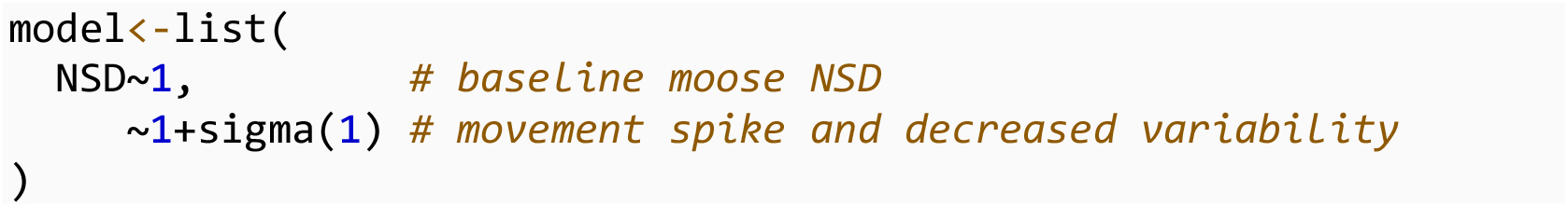

**Figure 2:**
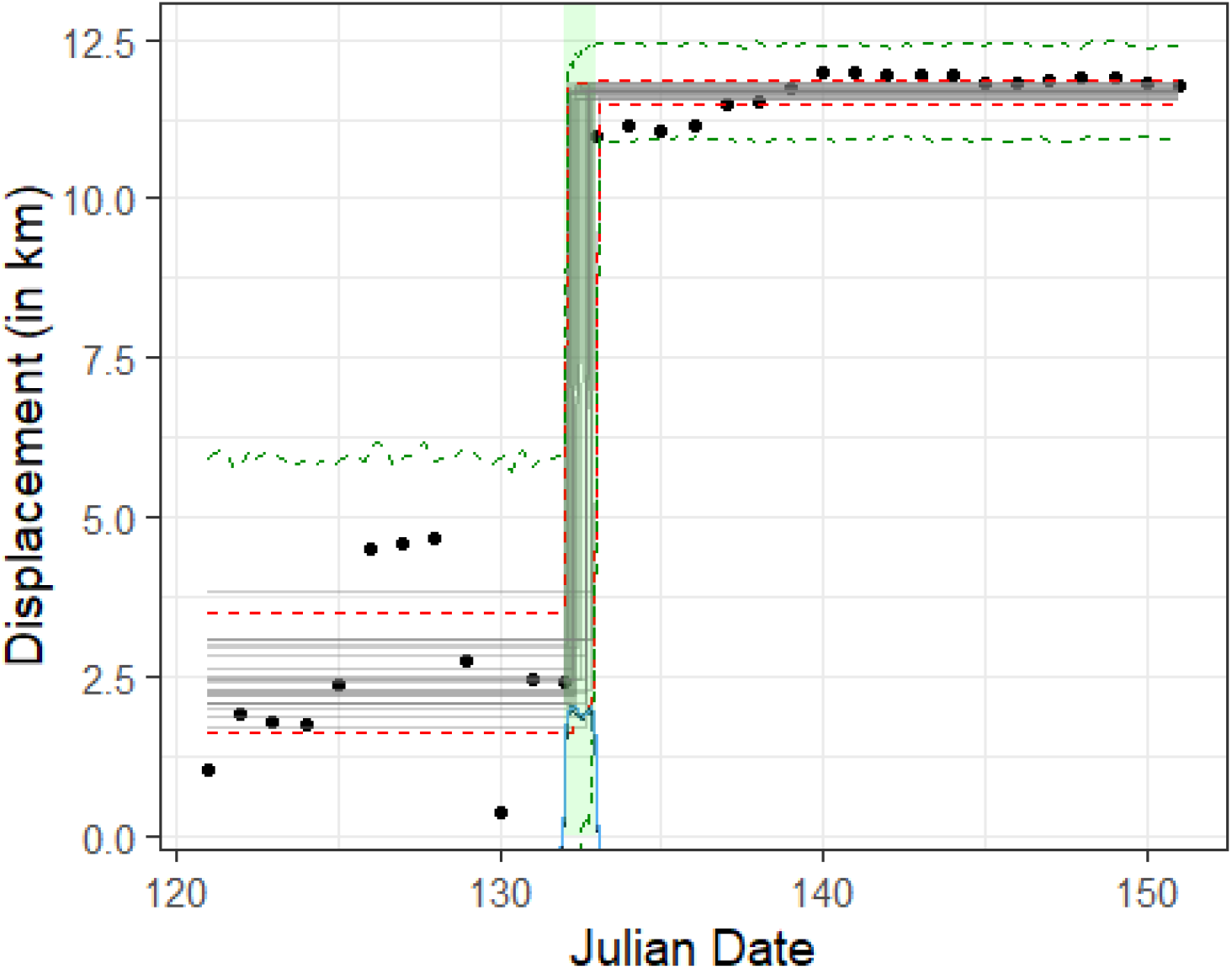
Displacement (distance from each location to the location at the start of the observation period) for a pregnant moose in Minnesota. The green area is the timeperiod identified by researchers as immediately preceding parturition. Grey lines represent 25 draws from the posterior distribution of the mean displacement. Red dotted lines depict 95% credible intervals for the mean displacement and green dotted lines depict 95% prediction intervals. The posterior distribution for the change point is shown in blue on the x-axis.

LOO-CV of this model versus one without a change in variance reflected that this model is a better fit to the data, thus illustrating the flexibility in adapting model syntax for each segment that corresponds to the biological situation. Prediction intervals illustrate that the addition of a change in variance for the second segment provides increased precision, therefore allowing the model syntax to closely mirror the biological situation of increased sedentary behavior (Figure 2).

Parturition events can be monitored in real time for studies involving the capture of neonates (Figure 2; Obermoller et al., 2019), or events can be identified post-hoc for retrospective analyses of ungulate breeding and fecundity (Bonar et al., 2018; Long et al., 2009). Previous studies have used piecewise regression to identify change points indicative of ungulate parturition based on movement, but implementation of the mcp package can provide additional analytical options (Berg et al., 2021).

Posterior predictive checks, a common method used to test goodness of fit, generate data from the fitted model by simulating from the posterior predictive distribution to visualize how well a model matches the observed values. Using the fitted model for moose movement, we demonstrate a graphical method to compare *y*, the observed response dataset, and *y*^*rep*^, replicated datasets simulated from the fitted model. (Figure 3). The two peaks at 3 km and 12 km correspond to the two intercepts fit by the model and overall, the observed data conform well to the replicated datasets.

**Figure 3:**
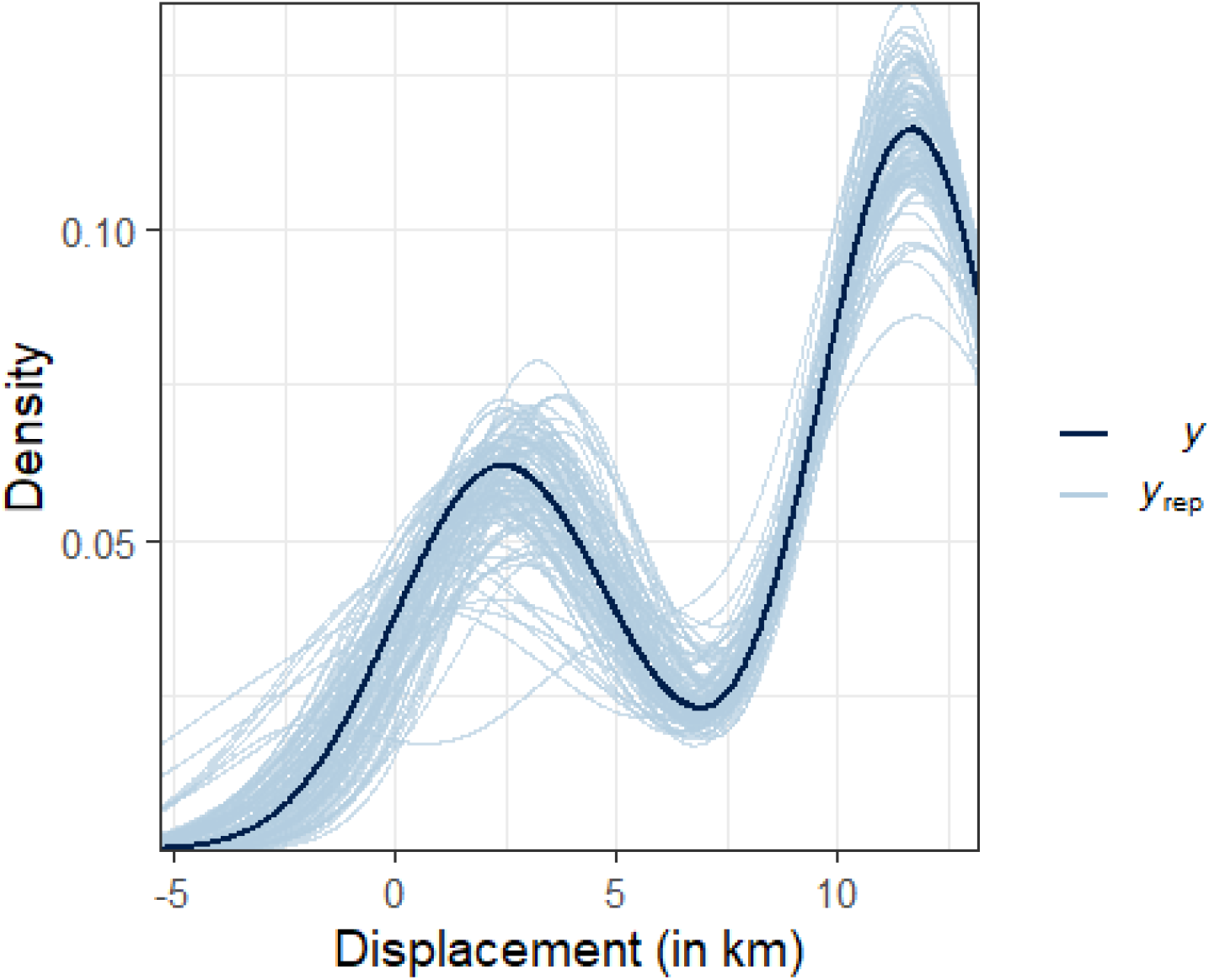
A kernel density posterior predictive check compares the distribution of observed outcomes (displacement in kilometers by a pregnant moose), shown in the black line, against 50 kernel densities of replicated datasets produced by the fitted model, each shown as a light blue line.

Prior field research can inform the ideal time window for change-point movement analysis and the scale of long-distance movement preceding parturition (allowing for a more informed prior).

### Physiological response to drone fly-over

Remotely piloted aircraft systems (hereafter drones) allow increased opportunity for remotely monitoring wildlife populations, and recent reductions in their cost have led to widespread adoption as an alternative to traditional aerial surveys using planes and helicopters (Watts et al., 2010). Drones instrumented with remote sensing technologies are now commonly used to estimate animal abundance in remote locations, collect fine-grain aerial imagery, and monitor poaching activities (Anderson & Gaston, 2013; Linchant et al., 2015; Mulero-Pázmány et al., 2017). The use of drones can decrease fieldwork costs and eliminate the need for hazardous fieldwork, and they can be coupled with computer vision methods to increase the quality and precision of data collection (Chabot & Francis, 2016; Hodgson et al., 2018; Seymour et al., 2017).

Despite the advantages of using drones for field data collection, their presence can influence behavior or illicit a physiological response in the study species, especially when they are flown at low altitudes (McEvoy et al., 2016; Mulero-Pázmány et al., 2017). Response to anthropogenic stimuli such as noise has been shown to have detrimental effects on wildlife species at both immediate (e.g., increased vigilance and decreased foraging behavior) and long-term scales (e.g., decreased reproduction and population declines: Shannon et al., 2016; Blickley & Patricelli, 2010; Senzaki et al., 2020). The response of animals to drones has focused on external responses, such as altered movement and behavior. Most research on internal physiological responses have been limited to quantifying levels of glucocorticoids (e.g., cortisol and corticosterone: Baker et al., 2013; Bennitt et al., 2019; Millspaugh & Washburn, 2004; Vas et al., 2015; Wasser et al., 2000).

More recently, biotelemetry data have been paired with physiological data, allowing for new insights into the response of animals to anthropogenic stimuli. For example, Ditmer et al. (2015) measured changes in movement and heart rate levels of black bears (*Ursus americanus*) affixed with GPS collars and internally-implanted cardiac biologgers during drone flights. Although drones rarely elicited a behavioral response in black bears, heart rate levels were strongly correlated with proximity of drones overhead (Figure 4).

**Figure 4:**
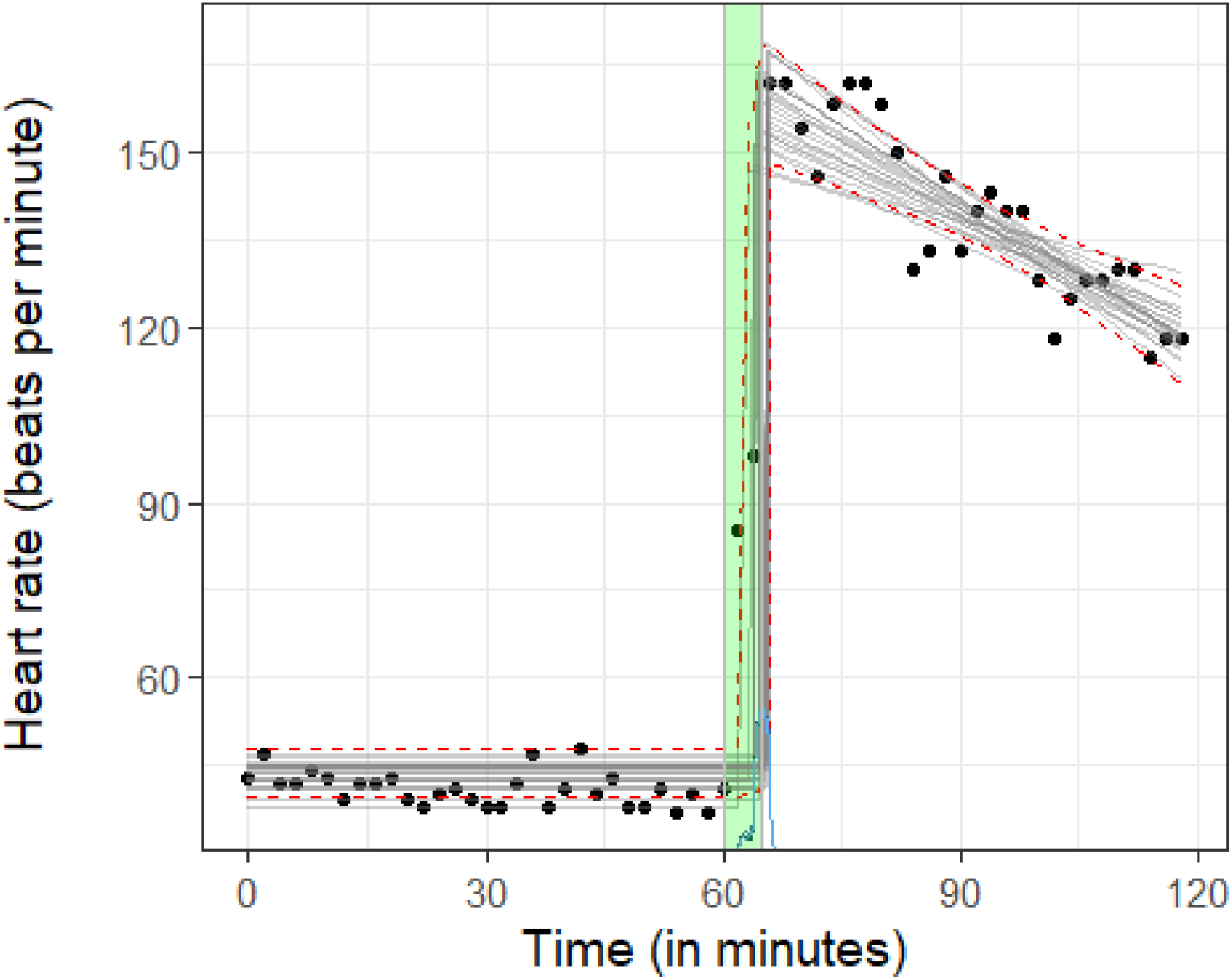
Heart rate of a black bear in Minnesota in relation to a drone flight. The green box indicates the duration of a drone flight. Grey lines represent 25 draws from the posterior distribution of the mean response. Red dotted lines depict the 95% credible intervals for the mean response. The posterior distribution for the change point is shown in blue on the x-axis.

Figure 4 shows the relationship between drone presence and black bear heart rate in beats per minute (bpm) before and after controlled flights (Ditmer et al., 2015).

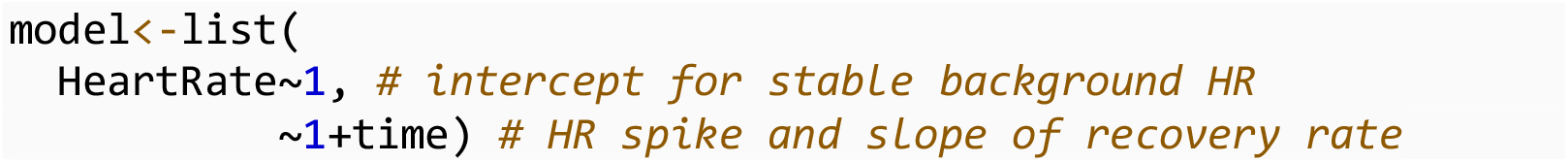

The initial segment prior to the change point includes an intercept capturing the baseline heart rate of the bear. The second segment contains an intercept and a disjoined slope term reflecting the heart rate gradually decreasing after the initial spike caused by the drone flight.

The flexibility of piecewise regression to accommodate different model structures for each segment allows for identification of the timing, magnitude, and acclimation rate of the stress response caused by the drone presence. For example, the change point, cp_1, is estimated to occur approximately 65 (95% CI=63-66) minutes into the observation period, the baseline heart rate, int_1, is estimated to increase by 116 bpm (int_2-int_1) once the drone appears, the heart rate then decreases by the slope of 0.78 bpm (minutes_index_2) and if the recovery rate is static, the black bear will return to baseline heart rate in approximately 2.5 hours ((int_2-int_1)/minutes_index_2=149 minutes). Lastly, sigma_1 captures the variability in heart rate about the overall trend.

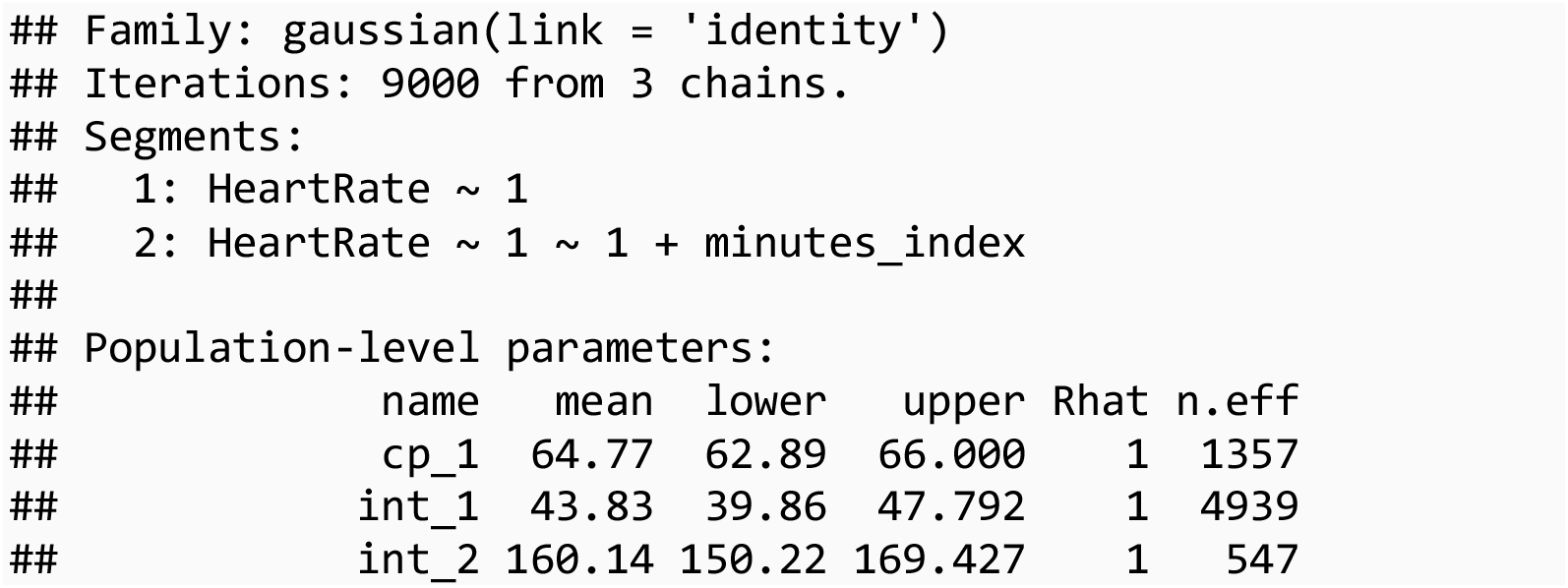
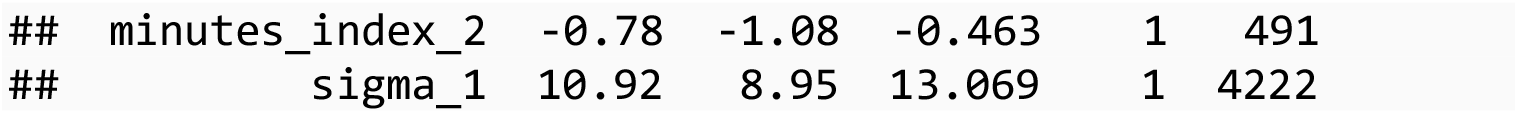

### Mortality signal from acceleration and temperature data

Accurate estimates of vital rates such as survival and fecundity are necessary for estimating population growth rates. Survival rates can be estimated using mark-recapture and mark-resight methods. However, for wide-ranging migratory species, individuals in subsequent years may not be resighted due to the combination of three separate possibilities: mortality, emigration, or missed detection by observers (Anders & Marshall, 2005). Because it is difficult to separate these components, biologists are often forced to estimate ‘apparent’ survival, as opposed to true survival (Lebreton et al., 1992). Additionally, mark-recapture and mark-resight methods do not provide information on specific times or locations of mortalities.

Many studies have illustrated the importance of evaluating vital rates over the entire annual cycle (Rushing et al., 2017; Sillett & Holmes, 2002). The miniaturization of GPS attachment devices and increased use of radiotracking now allows for a more direct accounting of survival during the breeding, migration, and overwintering seasons (Kays et al., 2015). Increased precision in vital rates during different seasons (and of different age classes) can better inform ecological studies considering life-history tradeoffs between survival and fecundity, especially concerning survival during migration periods (Buechley et al., 2021; Cheng et al., 2019; Flack et al., 2016).

Despite the advantages that GPS telemetry can offer to survival rate estimation, mechanical failure of transmitters can obscure whether the true fate was mortality or equipment failure. Some degree of failure is an unfortunate reality of most GPS tracking studies (Klaassen et al., 2014). Recovery of a GPS transmitter after a mortality event is often logistically unfeasible for species that migrate long distances. Therefore, a variety of methods have been employed to infer mortality versus transmitter failure from remotely obtained transmitter signals (Buechley et al., 2021; Sergio et al., 2019). The most common method applied is a visual assessment of whether locations appear to be stationary. However, this approach may lead to different conclusions depending on who makes the visual assessment (Koczur et al., 2017; Nygård et al., 2016; Rotics et al., 2017).

Sensor data such as battery voltage, temperature, and accelerometry are increasingly being used as diagnostic criteria for determination of mortality events versus transmitter failure (Burnside et al., 2016; Ely & Meixell, 2016; Hewson et al., 2016). Typically, mortalities coincide with shifts in the trend or variability in these data. Piecewise regression is therefore a quick and easy method that can be used to identify breakpoints that may be indicative of mortality. This approach can also be extended to monitor nest success with temperature loggers (Hartman & Oring, 2006; Sutti & Strong, 2014; Zangmeister et al., 2009). As part of a study of trumpeter swan movement ecology, we evaluated whether we could detect mortality using piecewise regression of Overall Dynamic Body Acceleration (ODBA) and temperature data (Figure 5; Wolfson, unpublished data).

**Figure 5:**
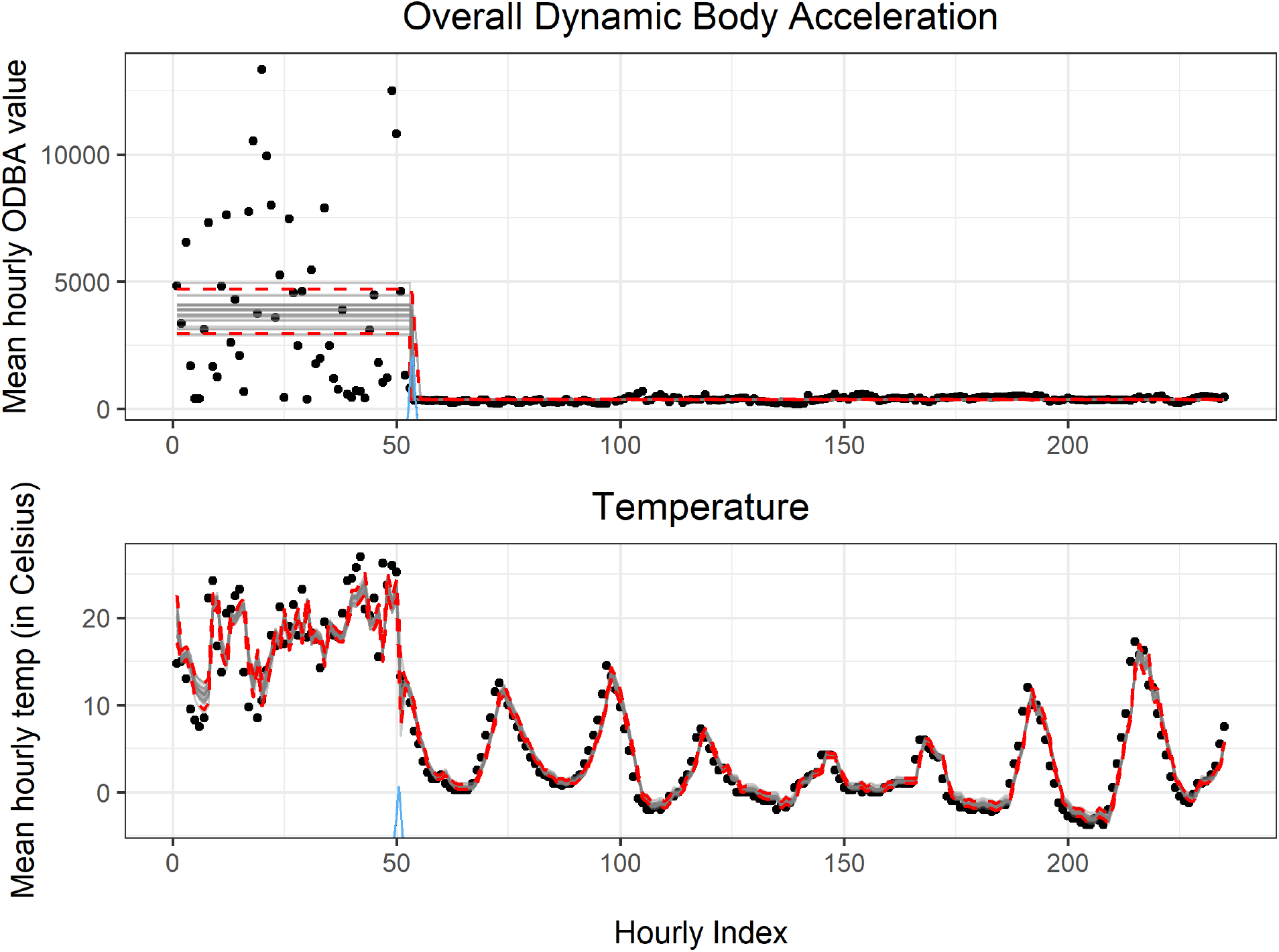
Time along the x-axis is an index since the start of the time series for Overall Dynamic Body Acceleration (ODBA) on the top figure, and temperature on the bottom figure. Grey lines show 25 draws from the posterior distribution, with 95% credible intervals for the mean response shown as red dotted lines. The posterior distribution for the change point is shown in blue on the x-axis.

The ODBA model includes a breakpoint separating two segments with differing means and variances.

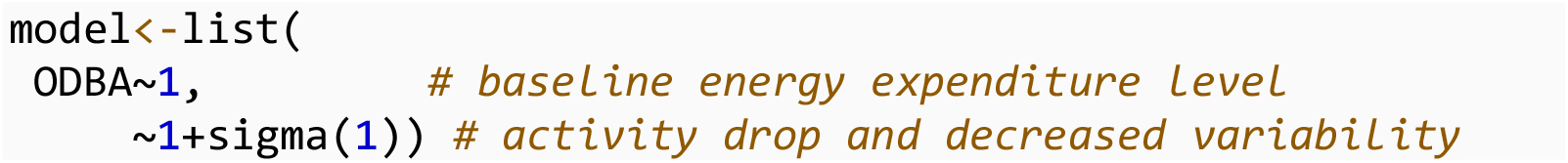

The temperature model has a breakpoint separating the first segment, which is modeled with a second-order autoregressive term, and the second segment, which is modeled with a first-order autoregressive term.

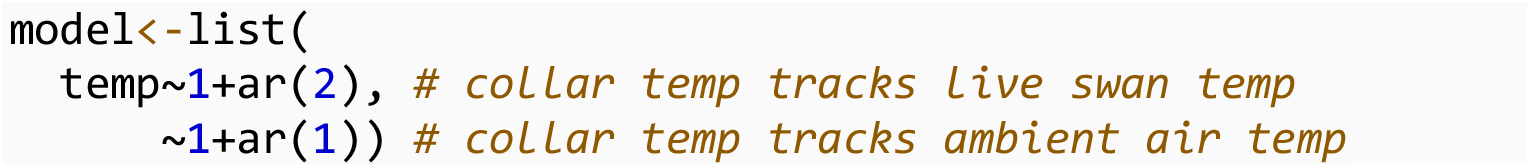

Both models showed a clear distinction at the change point representing mortality of the collared swan. There are advantages to being able to identify true mortality using multiple auxiliary data sources. Many transmitters are not equipped with tri-axial accelerometers, either to save costs or because accelerometer data is not a focus of the study. Additionally, deriving ODBA estimates from raw sensor data adds an additional analytical step to the process of estimating mortality.

### Migration phenology including stopovers

Seasonal migration allows species to optimize energetic budgets and undergo reproductive cycles while avoiding harsh environmental conditions and low food availability (Newton, 2010). Knowledge of migration phenology throughout the annual cycle inform management activities such as timing of water drawdowns and annual surveys to occur during peak migration, advance understanding of disease dynamics based on timing and overlap with other species, and reveal a species’ capacity to adapt their migratory timing in response to climate change to preserve optimal breeding conditions and peak food availability (Donnelly et al., 2019; Moller et al., 2008; Newman et al., 2009; O’Neal et al., 2012; Thurber et al., 2020).

Despite the importance of understanding migration phenology and common reporting of phenological information in movement-related wildlife analyses, a consistent methodology for determining departure and arrival dates has not yet emerged (Cerritelli et al., 2020; Soriano-Redondo et al., 2020). Often, there is no straightforward reproducible methodology, and authors provide vague descriptions of methods used to assign migration phenology to each individual (Shephard et al., 2015; Shimada et al., 2014). One of the simplest repeatable approaches is to segment phases of the migration cycle based on date ranges informed from prior information (Takekawa et al., 2010; Wolfson et al., 2017a).

Other approaches include using a spatial threshold, based on either the absolute distance from the capture origin, breeding territory, or last location, a spatio-temporal threshold, based on a certain distance moved within a period, or a crossing of a chosen latitude or landmark (Flack et al., 2016; Giunchi et al., 2019; Rotics et al., 2016). Although these approaches may yield useful estimates of migration phenology, often they rely on arbitrary criteria or choices that are either species-specific or highly subjective, and therefore hard to generalize to other study systems.

Model-based methods, including non-linear theoretical movement models fit to NSD, are also commonly used to estimate migration phenology (Börger & Fryxell, 2012; Bunnefeld et al., 2011; de Grissac et al., 2016; Spitz et al., 2017). The use of piecewise regression to segment an annual cycle based on NSD allows for a model-based approach, but is more flexible than typical NSD modeling *sensu* Bunnefeld et al. (2011) and can also provide additional information such as the timing and duration of stop-overs, which is information not attainable using traditional NSD-based methods.

We demonstrate the utility of piecewise regression in assessing movements during the annual cycle of a migratory bird using a NSD time-series of sandhill crane (Antigone canadensis) locations in North America (Figure 6; Wolfson et al., 2017b; Wolfson 2018). Although a basic three-intercept model would sufficiently discriminate the summer and winter periods, adding additional intercepts reveals each major staging area that the crane used.

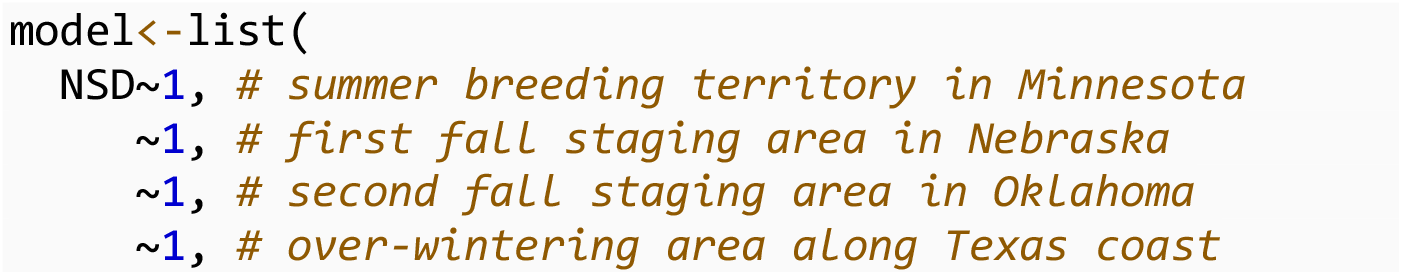

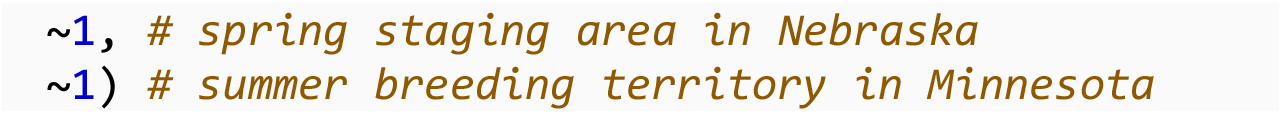

**Figure 6:**
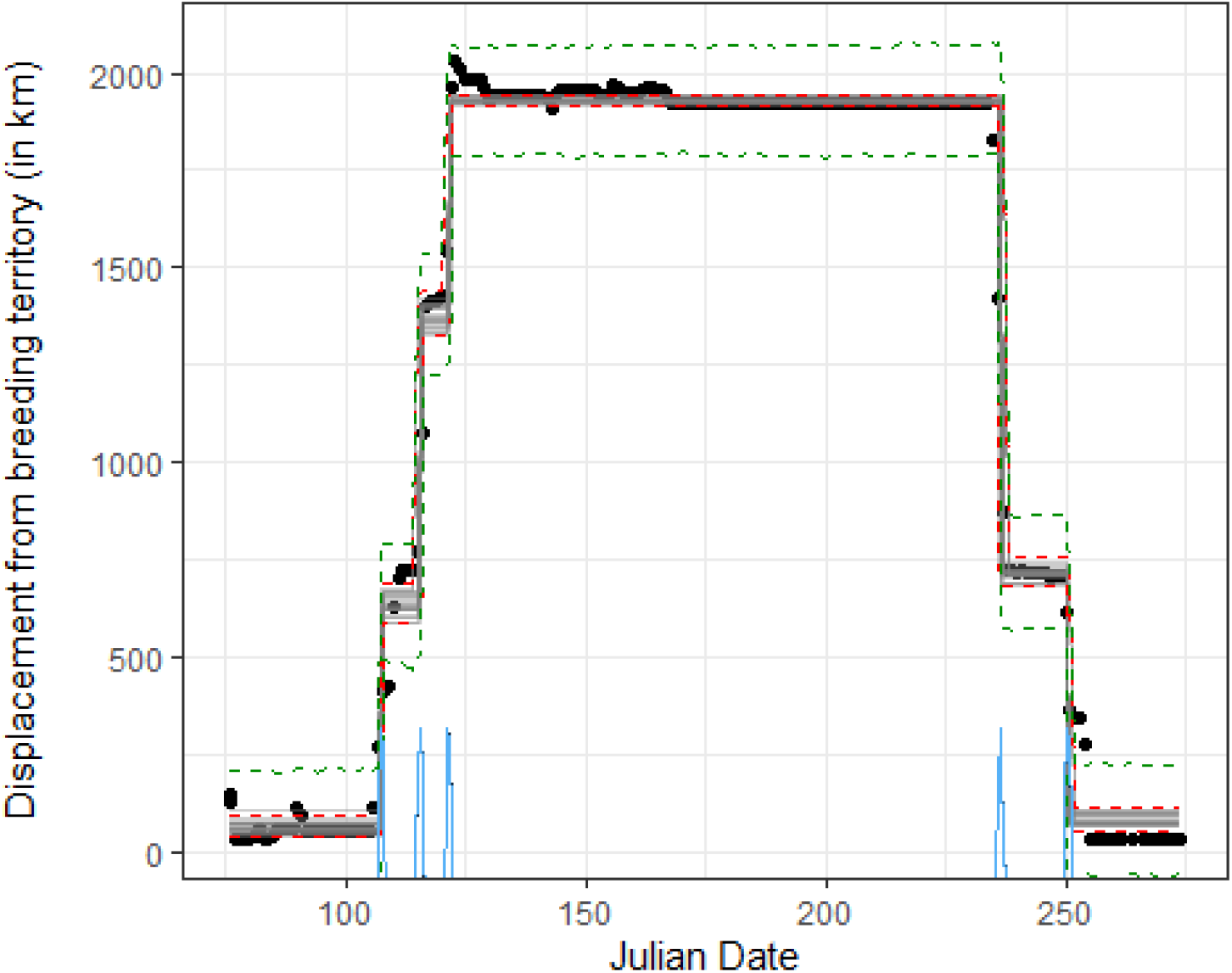
Time along the x-axis is an index since the start of the time series. The y-axis represents average daily displacement from the breeding territory in kilometers. Grey lines show 25 draws from the posterior distribution, with 95% credible intervals for the mean response shown as red dotted lines and 95% prediction intervals in green dotted lines. The posterior distribution for the change point is shown in blue on the x-axis.

The ideal number of stopover sites can be visually examined or empirically derived using cross-validation to compare multiple models with differing number of change points. Table 2 demonstrates that the model with six intercepts (representing two spring staging areas and one fall staging area) is the most well supported using ELPD as a model comparison metric.

**Table 2:** Leave-one-out Cross Validation was used to compute the Estimated Log Predictive Density (ELPD) for 7 models, each with an increasing number of intercepts. The highest ELPD for the model with 6 intercepts indicates that it has the highest predictive accuracy.

| Model Syntax | ELPD Difference | SE of Difference |
| --- | --- | --- |
| Six Intercepts | 0.00 | 0.00 |
| Five Intercepts | -0.48 | 0.83 |
| Seven Intercepts | -19.59 | 32.82 |
| Four Intercepts | -24.73 | 9.52 |
| Two Intercepts | -57.39 | 14.01 |
| Three Intercepts | -324.69 | 16.93 |
| One Intercept | -336.17 | 16.73 |

## Discussion

As we demonstrated with examples of birth, death, migration, and behavioral responses (both internal and external), piecewise regression is a flexible tool that can be used to identify a wide variety of biological phenomena across different taxa and data types. Piecewise regression, and change point detection in general, is most appropriate when the objective is to partition a dataset into heterogeneous segments that may correspond with different biological states. By partitioning a time-series, identification of a change point can identify the point in time when a biological response occurred.

The analysis of thresholds that dictate regime shifts is a common theme in studies that focus on community dynamics, trophic interactions, and ecosystem perturbations (Dodds et al., 2010; Gsell et al., 2016; Su et al., 2021). A common attribute of these areas is the presence of long-term time-series datasets. The volume of data in wildlife biotelemetry datasets is rapidly expanding and the degree to which researchers have fine-scale sensor data over time increasingly makes techniques such as piecewise regression an appropriate tool.

Although there are many R packages intended to detect change points, the mcp package represents the best combination to date of rigorous statistical methodology, flexibility, ease of use, and reproducibility (Erdman & Emerson, 2007; James & Matteson, 2014; Killick & Eckley, 2014; Lindeløv, 2020; Muggeo, 2008; Zeileis et al., 2002). The mcp package allows the user to specify a unique model syntax for each segment between break points, detect changes in mean, variance, and autocorrelation, and it also allows for robust inference using full posterior distributions for all parameters and change points.

Researchers with previous knowledge of the study system can directly incorporate this information when specifying parameter ranges and prior distributions. Although performance may slow with very large datasets, mcp includes parallel processing to increase computational efficiency. We hope that our worked examples will encourage other ecologists to consider the use of piecewise regression for identifying signals in telemetry and biologging data.

## Acknowledgements

We thank Victoria Drake, Nathan Cross, Jon Dachenhaus, David Fronczak, Gunnar Kramer, Emily Wells, Jeff Fox, Steve Cordts, Ed Zlonis, Bruce Davis, and Ciara McCarty, for assistance with fieldwork. We thank Mark Ditmer for access to black bear data and William Severud, Tyler Obermoller, Glenn DelGiudice, and Michelle Carstensen for access to moose data. Thanks to Amy Davis and Althea Archer for comments on a previous version of this manuscript. Funding for this project was provided by Minnesota Environmental and Natural Resources Trust Fund as recommended by the Legislative-Citizen Commission on Minnesota Resources (LCCMR) and the U.S. Geological Survey, Minnesota Cooperative Fish and Wildlife Research Unit. Any use of trade, firm, or product names is for descriptive purposes only and does not imply endorsement by the U.S. Government, the University of Minnesota, or the State of Minnesota. John Fieberg received partial support from the Minnesota Agricultural Experimental Station.

